# Loss of a glutaredoxin gene underlies parallel evolution of trichome pattern in *Antirrhinum*

**DOI:** 10.1101/518183

**Authors:** Ying Tan, Matthew Barnbrook, Yvette Wilson, Attila Molnár, Andrew Hudson

## Abstract

Most angiosperms produce trichomes--epidermal hairs that have protective or more specialised roles. In almost all species trichomes are multicellular and, in the majority, secretory. Despite the importance of multicellular trichomes for plant protection and as a source of high-value products, little is known about the mechanisms that control their development. Here we use natural variation between *Antirrhinum* (snapdragon) species to examine how trichome distribution is regulated and has evolved. We show that a single gene, *Hairy* (H), which is needed to repress trichome fate, underlies variation in trichome distribution patterns between all *Antirrhinum* species except one. *H* encodes an epidermis-specific glutaredoxin and trichome distribution within individual plants reflects the location of *H* expression. Gene phylogenies and functional tests suggest that *H* gained its trichome-repressing role late in eudicot evolution and that *Antirrhinum* species with widespread trichomes evolved multiple times from a largely bald ancestor though independent losses of H activity. We also find evidence for an evolutionary reversal involving a suppressor mutation, and for a pleiotropic effect of *H* that might constrain the evolution of trichome patterns.

## INTRODUCTION

Angiosperms produce epidermal hairs (trichomes) that are mostly associated with protection against herbivores or abiotic factors such as UV radiation (Bickford, 2016; Riddick & Simmons, 2014). In almost all species they are multicellular, and many have a similar structure of apical secretory cells supported by a stalk (Carlquist, 1961; Huchelmann *et al.*, 2017), which suggests homology and a single, ancient origin. Because trichomes protect plants and their secretions are the source of economically important compounds, including pharmaceuticals and flavours (Schilmiller *et al.*, 2008), regulation of their development is a target for crop improvement and biotechnology (Glas *et al.* 2012; Huchelmann *et al.*, 2017; Schilmiller *et al.*, 2008).

Although associated with beneficial roles in protection, trichomes often differ in distribution within individual plants (e.g., Evans *et al.*, 1994; Telfer *et al.*, 1997)) or between close relatives (e.g., Bloomer *et al.*, 2012). This suggests either that there are fitness costs of trichomes that trade off against protection or that changes in trichome distribution are developmentally constrained. These possibilities are difficult to distinguish for multicellular trichomes because control of their development is only poorly understood. Their formation in diverse dicots is known to require an HD-Zip IV transcription factor, consistent with the homology of multicellular trichomes (Vernoud *et al.*, 2009; Walford *et al.*, 2011; Wang *et al.*, 2016; Yan *et al.*, 2017; Yang *et al.*, 2011; Zhu *et al.*, 2018). There is also circumstantial evidence for a conserved role for MIXTA-like MYB transcription factors, because one family member (GhMYB25) promotes formation of multicellular leaf trichomes in cotton (Walford *et al.*, 2012), while other members from *Antirrhinum* can induce ectopic multicellular trichomes when mis-expressed in tobacco (e.g., Glover *et al.*, 1998). It appears clear, however, that the mechanism regulating multicellular trichomes is not shared with the unicellular trichomes of Arabidopsis (reviewed by Pesch & Hulskamp, 2009), suggesting parallel evolution of the two trichome types. For example, both require an HD-ZIP IV gene, though these were recruited from different paralogues that diverged before the origin of seed plants (Chen *et al.*, 2017; Zalewski *et al.*, 2013), probably from an ancestor that was already expressed in the epidermis (Javelle *et al.*, 2011).

One way to reveal more of the mechanism controlling multicellular trichomes, and of the constraints on its evolution, is to exploit natural variation. Here we apply this approach using the genus *Antirrhinum* (snapdragons), which has ~25 species differing in multicellular trichome morphology and the pattern of trichome distribution within plants. We show that a single gene, *Hairy*, accounts for a restricted distribution of trichomes in species that are largely bald. *Hairy*, which encodes a glutaredoxin, is needed to suppress trichomes fate and is expressed in epidermal cells in response to developmental phase. Combined phylogenetic and functional analysis suggests that *Hairy* was recruited to suppress trichome formation late in the evolutionary history of eudicots and that independent losses of Hairy activity were involved in parallel evolution of hairy *Antirrhinum* species.

## RESULTS

*Antirrhinum* species are divided into three morphological subsections that correlate with ecology–small, prostrate alpines in subsection *Kickxiella* and large, upright competitors in subsections *Antirrhinum* and *Streptosepalum* (Rothmaler, 1956; Sutton, 1988; Wilson & Hudson, 2011). While subsections largely reflect evolutionary relationships, there is evidence for gene flow between geographic neighbours and for parallel evolution of *Kickxiella-like* morphologies in two species within subsections *Antirrhinum* and *Streptosepalum*, although it unclear whether this involved introgression from the *Kickxiella* lineage, sorting of ancestral variation, or mutation *de novo* (Sutton, 1988; Wilson & Hudson, 2011).

Morphological subsections of *Antirrhinum* differ in the pattern of trichome distribution. All species produce glandular multicellular trichomes from leaves and internodes below metamer 4 (m4, where cotyledons and the internode above them are m1; Fig. 1c, d). Species in subsection *Kickxiella* remain densely hairy throughout their development, although species differ in trichome morphology from m4 (Fig. 1g-j). In contrast, leaf blades and stems above m4 are bald in subsections *Antirrhinum* and *Streptosepalum*, though glandular trichomes are retained on adaxial leaf midribs (Fig. 1b, f) and trichome production resumes in the inflorescence (Fig. 1a, d). Two aspects of trichome formation are therefore related to developmental phase: trichome morphology changes from around m4 in *Kickxiella* species while leaf blades and stems become bald from this point in subsections *Antirrhinum* and *Streptosepalum.* We found a few populations of four species in subsections *Antirrhinum* and *Streptosepalum* that were polymorphic for bald and hairy plants, possibly as a result of hybridisation with subsection *Kickxiella* (Sutton, 1988; Wilson & Hudson, 2011).

**Figure 1.**
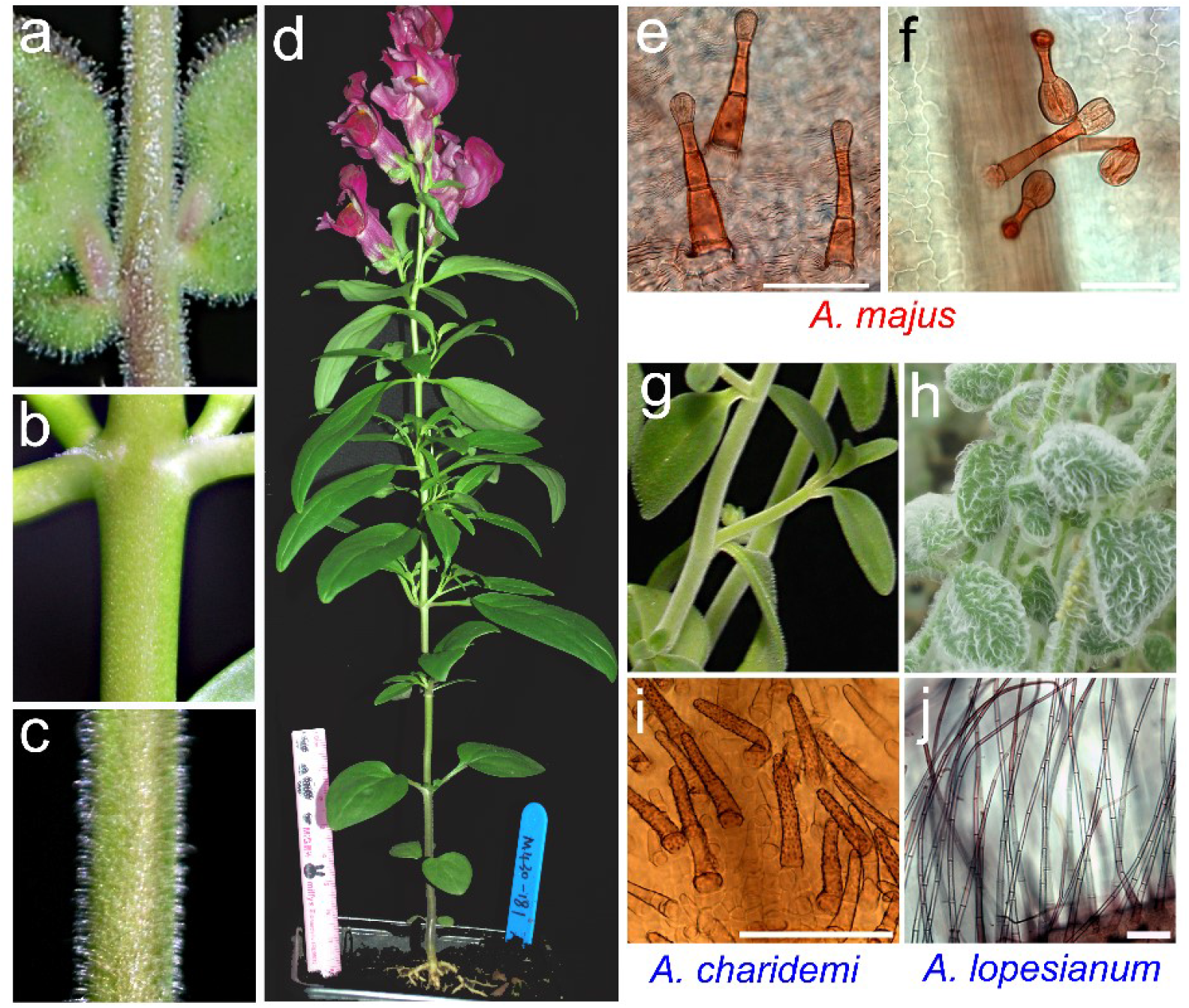
Variation in *Antirrhinum* trichome distribution and morphology. (a-f) *A. majus* (subsection *Antirrhinum)*, (a) inflorescence, (b) m5 stem and node, (c) m2 stem, (d) all aerial parts, (e) short glandular trichomes on m2 leaf blades and (f) the adaxial midrib of m5 leaves. (g-j) m5 leaves of two species in subsection *Kickxiella: A. charidemi* (g, i) with short glandless trichomes and *A. lopesianum* (h, j) with long glandular trichomes. Scale bars 0.1 mm.

To investigate the genetic basis for variation in trichome distribution, we first crossed hairy *A. charidemi* (subsection *Kickxiella;* Fig. 1g, i) with bald *A. majus* (subsection *Antirrhinum*, Fig. 1a-f). F1 progeny had the bald phenotype of the *A. majus* parent, suggesting that it carries a dominant inhibitor of trichome fate. Hairy plants occurred in the F2 at a frequency of ~6% (18 among 285) and in a genome scan were all found to be homozygous for the *A. charidemi CYC* allele, linked to the self-incompatibility (*S*) locus that prevents an active *S* allele, as from *A. charidemi* (*S^c^*), becoming homozygous (Schwarz-Sommer *et al.*, 2003). *A. majus*, in contrast, carries an inactive *s* allele (Qiao *et al.*, 2004). Therefore the inheritance of baldness can be explained by a single locus, *Hairy* (*H*), at which the *A. majus* allele, *H^m^*, is a dominant suppressor of trichomes and hairy F2 plants (*h^c^/h^c^* homozygotes) occur only when *h^c^* is uncoupled from *S^c^* by recombination, which occurred in ~13% of pollen (Supplemental Information).

We then screened a population of near-isogenic lines (NILs), produced from the F1 hybrid by repeated back-crossing to *A. majus*, to identify a NIL that remained heterozygous at *H* but did not carry *S^c^*. It produced ~25% hairy progeny after self-pollination, supporting the view that the single repressor locus, H, is responsible for differences in hair distribution between the parents. The NILs revealed two further aspects of H function. Firstly, hairy progeny made glandular trichomes throughout vegetative development, rather than the short glandless trichomes of *A. charidemi*, implying that genes other than *H* are responsible for differences in trichome morphology (Supplemental Fig. S1). Secondly, bald and hairy siblings did not differ significantly for other traits related to developmental phase, such as flowering time, suggesting that *H* responds to underlying developmental phase information rather than contributing to it (Supplemental Table S1).

The *H* locus was further mapped by sequencing DNA pools from hairy or bald offspring of the NIL. Mapping reads onto a draft of the parental *A. majus* genome showed the introgression from *A. charidemi* in the NIL extended over~ 4.4 Mb and that *H* was located within a region of ~104 kb in which only *A. charidemi* SNPs were detected in the hairy (*h^c^/h^c^*) pool (Fig. 2a, b).

**Figure 2.**
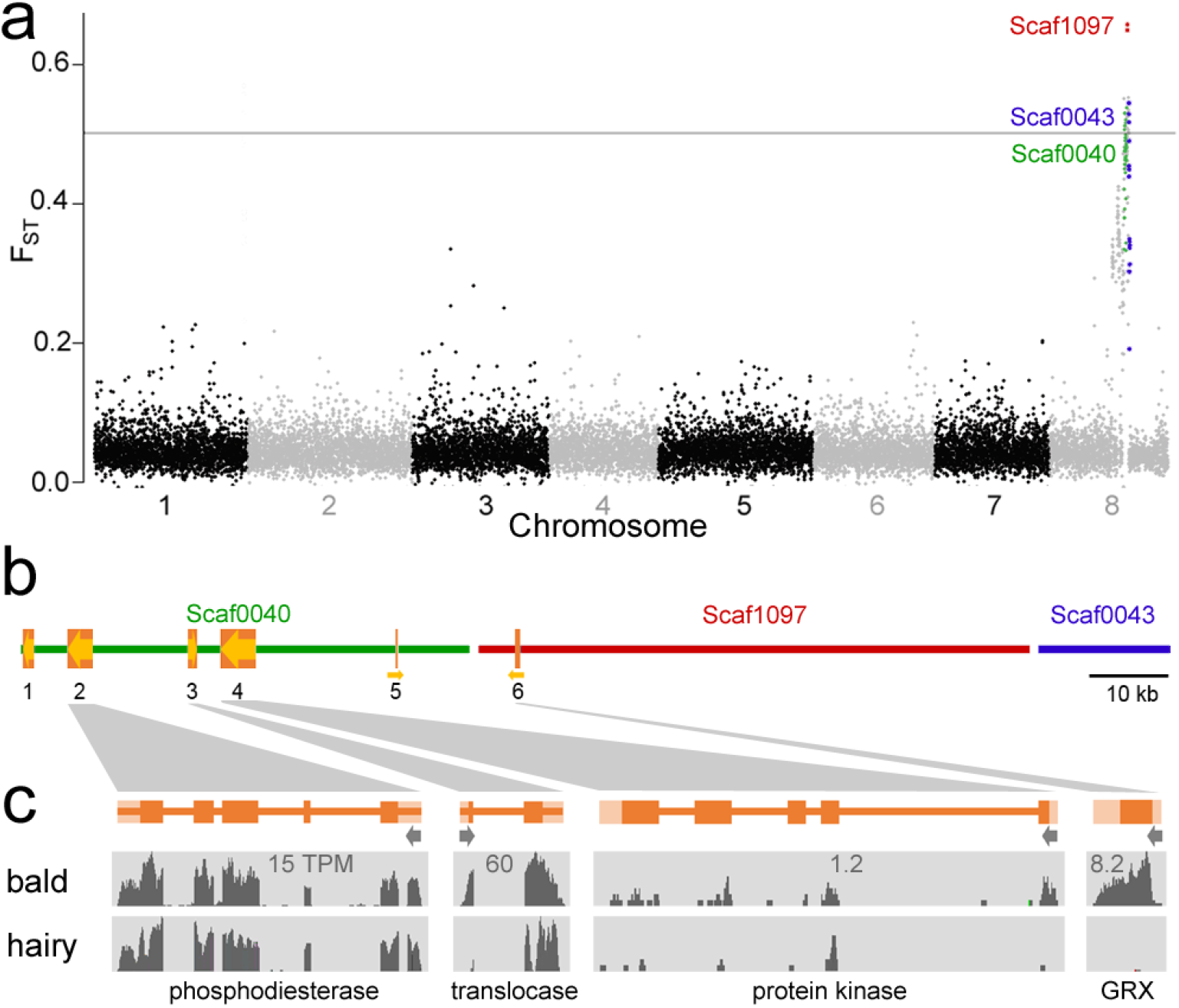
Mapping the *Hairy* locus. a) a scan of *F_ST_* between pools of hairy or bald progeny from an *h^c^/H^m^* near-isogenic line, shown in 10 kb bins with sequence scaffold order inferred from recombination maps. Around two-thirds of plants in the bald pool will be *h^c^/H^m^* heterozygotes, giving an expected *F_ST_* of ~0.5 for a marker at *Hairy.* b) The closest recombination breakpoints in hairy (h^c^/h^c^) progeny delimit *Hairy* to a region spanning three sequence scaffolds. Orange boxes show genes; the relative orientation of Scaffold 1097 could not be determined. c) Mapping of RNA-seq reads from bald and hairy phenotypes, with transcripts per million reads (TPM) values shown for the bald phenotype.

RNA-seq of bald and hairy NILs identified four genes within the target region that were expressed in vegetative apices above m4. *GRX8b* encoded a glutaredoxin in the land plant-specific CC-clade (Supplemental Information Fig. S2) and its RNA was present in bald apices but undetectable in hairy, suggesting that *H* encoded the GRX and that *h^c^* is a null mutation (Fig. 2c). A similar sequence *(GRX8a)* was found by homology in the target region, but was unlikely to contribute to H function because its expression could not be detected in any aerial tissue (Supplemental Fig. S3). The other genes in the interval showed similar levels of expression in bald and hairy phenotypes.

If *GRX8b* is H, then reducing its activity by virus-induced gene silencing (ViGS) (Liu *et al.*, 2002) should cause a hairy phenotype because *H* is a dominant suppressor of trichomes. *A. majus* infected with viruses carrying part of the *Phytoene Desaturase (PDS)* gene reported ViGS as tissue bleaching (Fig. 3a). Whereas reducing PDS activity alone had no effect on trichome production, adding either the coding region or the 3’UTR of the candidate *GRX8b* to the virus allowed trichomes to form from bleached leaves and stems above m4 (Fig. 3b-d, Supplemental Fig. S4), supporting the view that *GRX8b* is needed for H activity. ViGS appeared to be specific for the H-linked *GRX8b*, because the most similar expressed paralogue, *GRX6c*, was unaffected (Fig. 3f). Therefore *H* is very likely to be *GRX8b.* In contrast to H, ViGS with *GRX6c* did not alter trichome development, suggesting that this gene has a different role (Supplemental Fig. S4).

**Figure 3.**
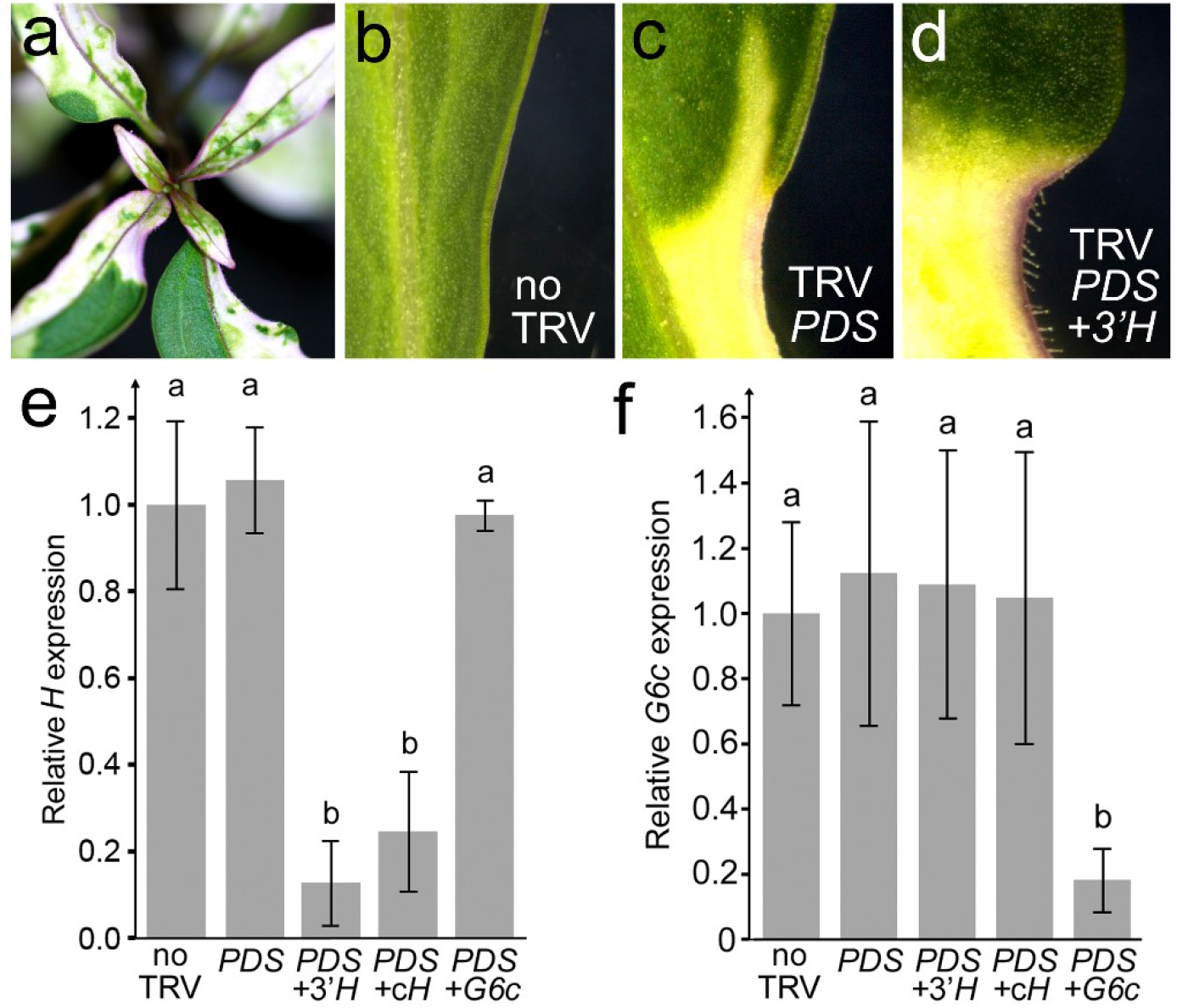
Hairy-linked GRX activity is required to suppress trichome formation. a) Tissues bleaching in *A. majus* following ViGS of *Phytoene desaturase (PDS)* with TRV carrying *PDS* and *Hairy* sequences. b-d) ViGS with part of the 3’-UTR of *Hairy* and *PDS* allows trichome development from bleached areas of stems and leaves at metamer 5 and above, whereas silencing *PDS* alone (b) has no effect on trichomes. e) Expression of the Hairy-linked *GRX8b* gene, but not its paralogue *GRX6c (G6c)*, is reduced by virus carrying either the Hairy-linked *GRX* coding region (cH) or it 3’-UTR (*3’H*), while *Hairy-GRX* expression is not reduced significantly by ViGS of *G6c* (f). Each value is the mean (±SEM) from three different plants. Means different with *p*≤0.01 are shown with different letters.

*H* is needed to suppress trichome formation only between m4 and the inflorescence. Consistent with this spatially restricted activity, we detected expression of the *H^m^* allele only in apices producing bald leaves and internodes (Fig. 4a). Within these, in situ hybridisation showed *H* RNA confined to the epidermal cells of leaf primordia, from around the developmental stage at which trichomes begin to form in hairy plants (Fig. 4b). Expression was not detected in epidermal cells of the adaxial midrib, which produce trichomes in all genotypes (Fig. 4c). Therefore the pattern of *H* activity and the distribution of trichomes appear to reflect *H* RNA expression. In contrast, mRNA from the *GRX6c* paralog was present in all aerial tissues tested and restricted to internal cells (Fig. 4e-h), supporting a role that is not related to trichome development. However, like H, its expression changed with developmental phase (Fig. 4e), suggesting that its common ancestor with *H* might have been phase-regulated.

**Figure 4.**
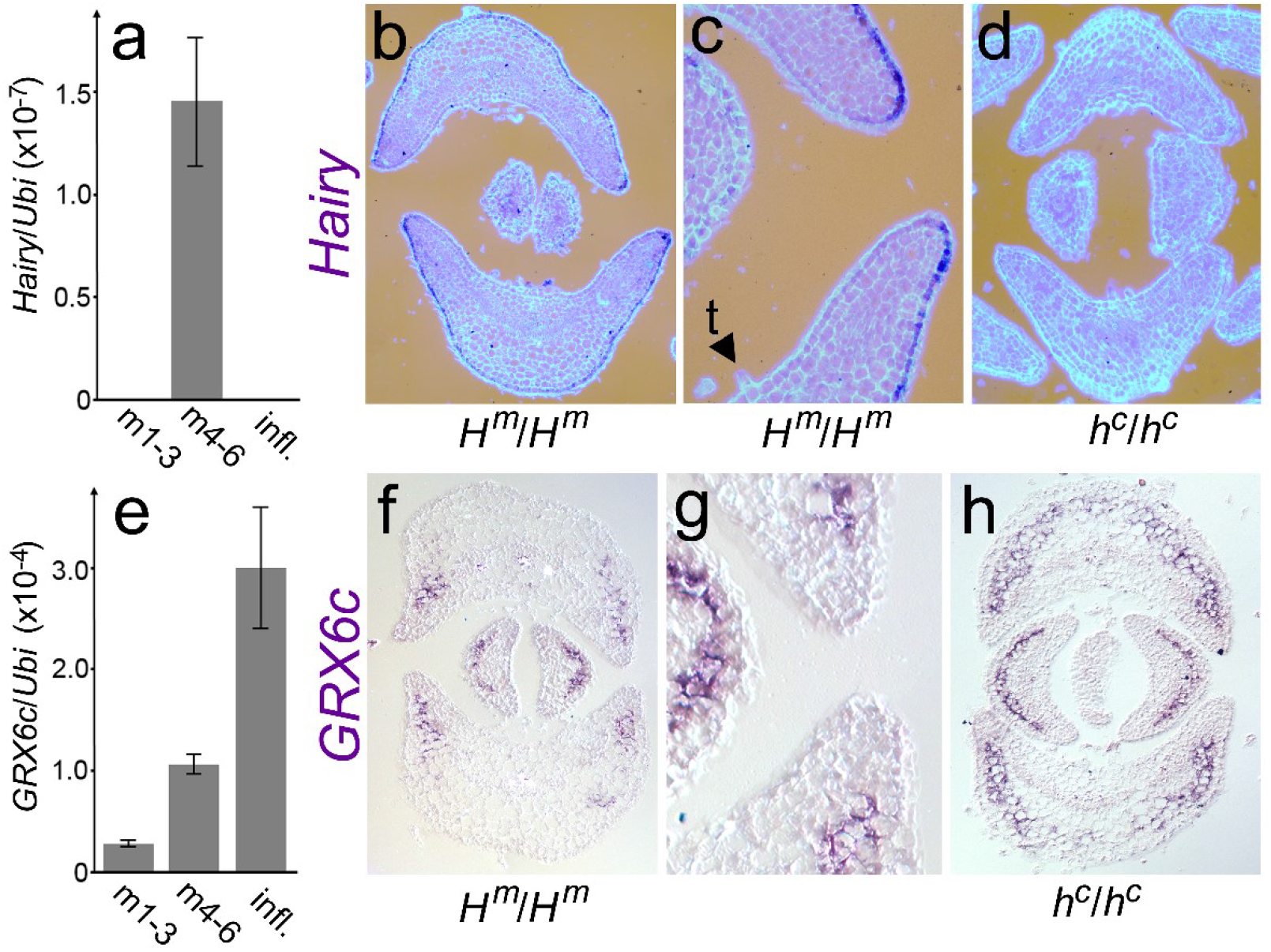
*Hairy* is expressed in the epidermis of bald leaves. a) Quantitative RT-PCR of *Hairy* mRNA, relative to *Ubiquitin (Ubi)*, in vegetative apices at different stages of development. b-c) In situ detection of *Hairy* RNA in transverse sections of bald *(H^m^/H^m^)* or hairy (h^c^/h^c^) apices. The younger, central pair of leaf primordia are at metamer 6, t indicates a trichome formed from the adaxial midrib. e) mRNA levels of the *Hairy* paralogue *GRX6c* and f-h) in situ hybridisation with a *GRX6c* probe.

To understand the origin of H activity, we identified the sequences most similar to *H* from two other members of the tribe *Antirrhineae (Misopates orontium* and *Chaenorrhinum origanifolium)* and in genome sequences from more distant eudicot lineages (Ogutcen & Vamosi, 2016; Zeng *et al.*, 2017). Phylogenetic analysis placed *H* in a well-supported clade specific to the *Antirrhineae*, and clustered *Antirrhinum H* with the *M. orontium* and *C. origanifolium* sequences, implying their orthology (Fig. 5a). To test further the function of the *M. orontium* orthologue *(MoHairy)* we reduced its expression by ViGS. Uninfected plants become almost bald at higher vegetative metamers (Fig. 6b) whereas stems experiencing ViGS were able to produce dense trichomes throughout development (Fig. 6c, d), revealing a conserved role in trichome suppression for *H* and *MoHairy.* The more basal *Antirrhinum* genes in the same clade do not appear to regulate trichomes--GRX6c is expressed internally and ViGS with this gene does not affect trichome development, while *GRX8a* is not expressed in shoots. Therefore *H* seems likely to have acquired its trichome repressing role relatively late in the evolutionary history of angiosperms, within the Lamiales and after divergence of the lineages leading to Mimulus and the *Antirrhineae* but before the *Antirrhinum–Misopates* split. This is consistent with H being co-opted to repress a more ancient mechanism that promotes multicellular trichome fate.

**Figure 5.**
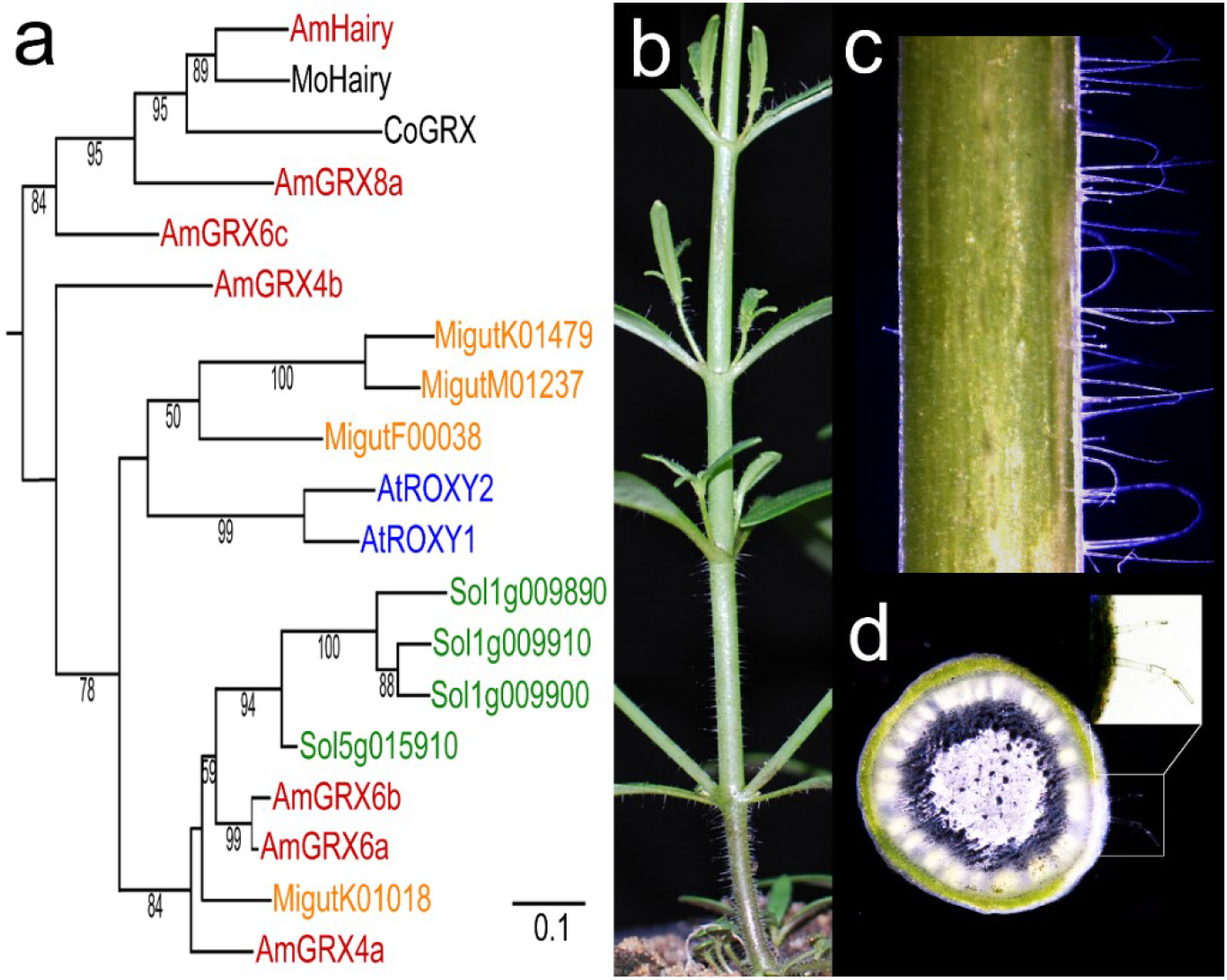
Origin of the *Hairy* gene. a) Maximum-likelihood tree of *A. majus* (Am) Hairy protein and the most similar sequences from *A. majus* (red), *Misopates orontium* (MoHairy), *Chaenorrhinum origanifolium* (CoGRX), Mimulus (orange), tomato (green) and *A. thaliana* (blue). *A. majus GRX* genes are numbered according to their chromosome location. Bootstrap support ≥50% is shown. The tree is rooted on a sister clade (Supplemental Fig. S7). b) *M. orontium* stems become bald from node 3. c) ViGS of *MoHairy* and *PDS* genes. Only the right-hand part of the stem shows silencing, revealed by tissue bleaching in a transverse section (d), with details of ectopic trichomes in bright-field illumination inset.

We then examined whether *H* could account for all variation in trichome distribution within the genus *Antirrhinum.* Except for a few polymorphic populations, the hairy phenotype is a characteristic (synapomorphy) of subsection *Kickxiella* (Wilson & Hudson, 2011). The simplest evolutionary scenario is therefore that the ancestral *Antirrhinum* had H activity and was bald, consistent with the conserved role of *H* in the *M. orontium* outgroup, and that H activity was retained in subsections *Antirrhinum* and *Streptosepalum* but lost once at the base of the main *Kickxiella* lineage.

To test these hypotheses, we first crossed the heterozygous *h^c^/H^m^* NIL to different species. Each cross to a hairy *Kickxiella* species, or to a hairy member of a polymorphic population, produced ~50% hairy progeny, suggesting that all hairy taxa lacked H activity (Supplemental Information Table S2). Conversely all progeny of crosses to bald species, with one exception, were bald, consistent with active *H* alleles in bald species. The exception, *A. siculum*, produced hairy progeny when crossed with a hairy NIL (h/h), and bald progeny from the cross to *A. majus* (H/H), suggesting that it is an *h* mutant but is bald because it also lacks activity of a gene required for trichome formation (Supplemental Information). This second-site mutation could have been involved in an evolutionary reversal from the hairy to bald state.

We then related trichome distribution within the genus *Antirrhinum* to *H* haplotypes. The whole locus appeared to have been deleted in a closely-related group of endemic *Kickxiella* species (Supplemental Table S3, Fig. S5). Sequences from the remaining *Kickxiella* species, with the exceptions of *A. hispanicum* and *A. grossi*, formed a single clade, consistent with a single *h* loss-of-function mutation in their common ancestor (Fig. 6). None of the alleles tested from this clade produced detectable RNA (open circles in Figure 6), suggesting that the ancestral mutation abolished expression and that additional substitutions and deletions accumulated in the absence of purifying selection. This might also be true of the deletion of *H* in the endemic *Kickxiella* species. The clade also contained nonfunctional *h* alleles from polymorphic populations of *A. latifolium* and *A. graniticum*, supporting their transfer from subsection *Kickxiella* by hybridisation. Both species are self-incompatible, and frequency-dependent selection favouring rare *Kickxiella S* alleles might have helped linked *h* alleles to increase in frequency after hybridisation (Wright 1939). A single *Kickxiella* haplotype was also found in all *A. siculum* samples, corroborating genetic evidence for *A. siculum* being an *h* mutant.

**Figure 6.**
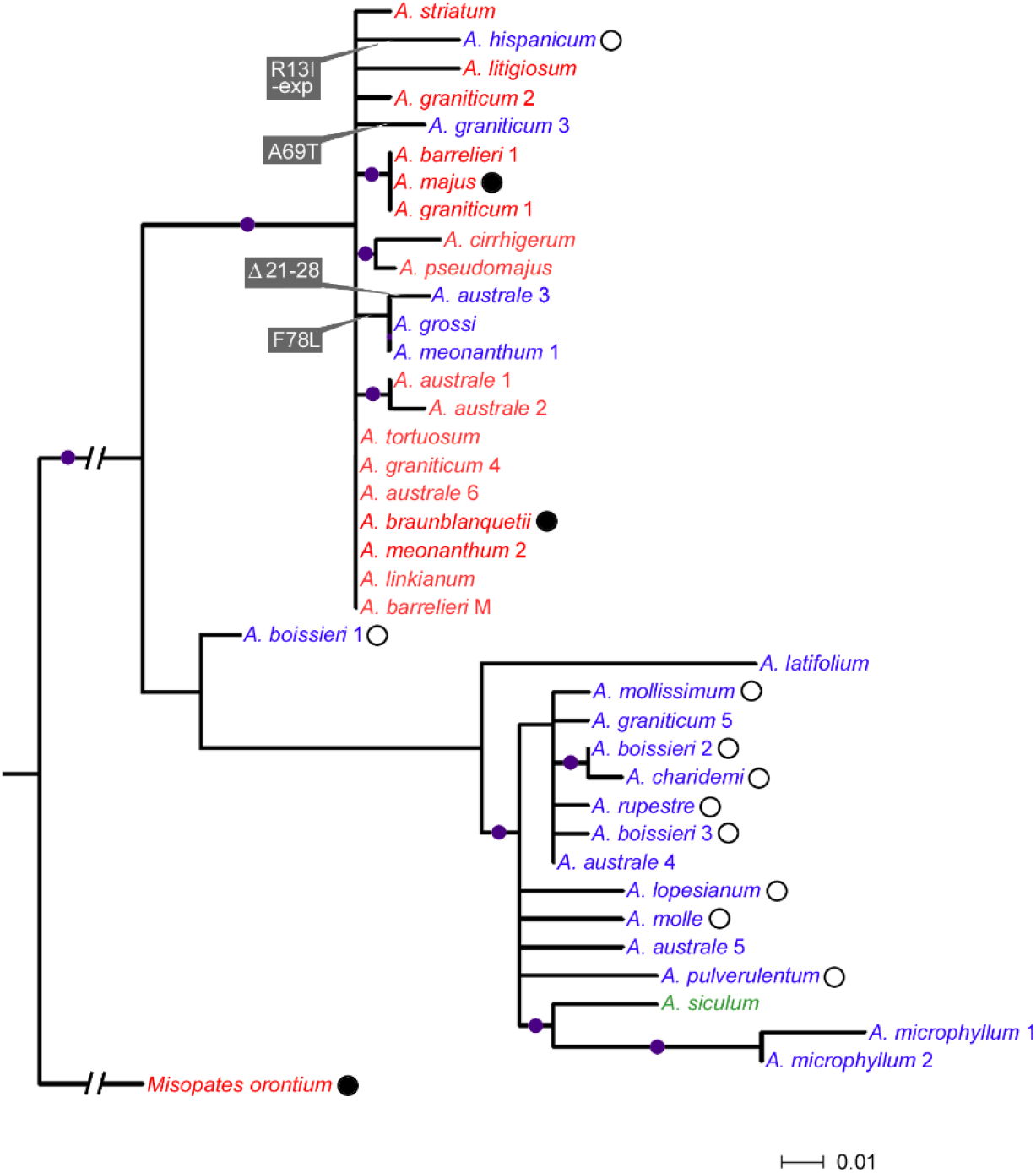
Evolution of the *Hairy* gene within *Antirrhinum*. A maximum-likelihood tree of the *H* open reading frame. Functional *H* alleles, inferred from phenotypes and allelism tests, are shown in red and non-functional *h* alleles in blue. Expressed alleles are marked with filled circles, those for which no expression could be detected with open circles. Nodes recovered in ≥50% of bootstrap replicates are marked with dots. More than one allele was identified in some species; the distribution of haplotypes between species and populations is given in Supplemental Table S3. Non-synonymous substitutions that can explain independent losses of function are in grey boxes. The *A. australe* 3 allele shares the F78L substitution with two other non-functional *h* haplotypes, but also has a unique deletion of eight amino acids, shown by an artificially lengthened terminal branch that is not to scale. The *A. hispanicum* allele has a unique R13I substitution and also appears to have lost expression.

The active *H* alleles from bald species in subsections *Antirrhinum* and *Streptosepalum* formed a separate clade, with relatively little diversity and extensive sharing of haplotypes between species (Fig. 6, Supplemental Fig. S9). However, this clade also included five inactive *h* alleles, as inferred from allelism tests, each differing from its ancestral active allele by at least one non-synonymous substitution that could explain loss-of-function. They included *h* alleles from *A. australe* and *A. meonanthum*, which share an amino acid substitution (F78L), suggesting that an independent mutation had contributed to trichome polymorphism in populations of these two species, and from *A. hispanicum* and *A. grossi*, consistent with parallel evolution of their *Kickxiella-like* morphologies having involved *de novo* mutations, rather than introgression from *Kickxiella.* The hairy character therefore appears to have evolved at least four times independently from the ancestral bald state of *Antirrhinum* (Fig. 6) and possibly reverted once to baldness in the ancestor of *A. siculum*, via a second-site suppressor mutation.

Trichomes are associated with beneficial roles, raising the questions of how the bald character and H function arose. Similarly, the reversal from bald to hairy appears relatively unconstrained because ViGS revealed that it requires only loss of H activity, yet many species remain bald. To investigate whether *H* has pleiotropic effects that might constrain its evolution, we compared development of bald and hairy progeny of the NIL. Hairy (h^c^/h^c^) plants produced significantly larger leaves from m6 onwards compared to their bald (*H^m^*/*H*^m^) siblings (Fig. 7). Although a gene closely linked to *H* could explain this difference, it was detected only in leaves that express H. This supports a pleiotropic effect of *H* in suppressing both trichome formation and leaf growth that might expose the locus to selection acting in different ways.

**Figure 7.**
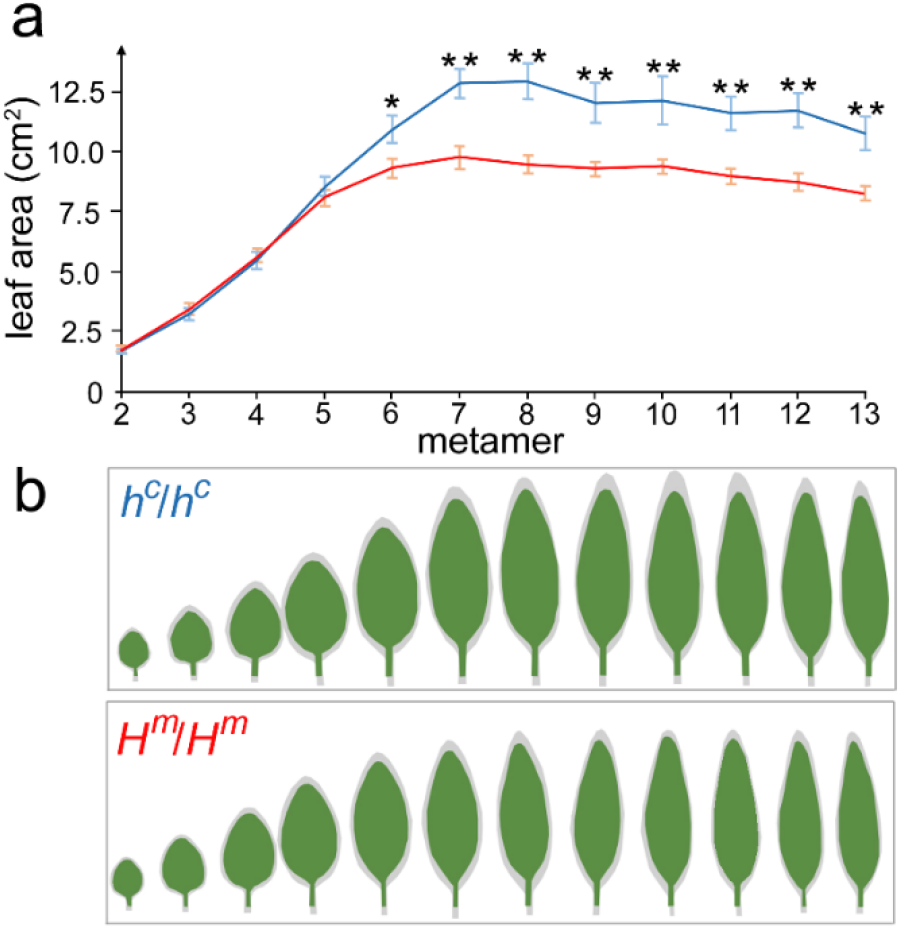
The *H* locus affects leaf area. a) A comparison of leaf areas for hairy (h^c^/h^c^, blue) and bald (H^m^/H^m^ red) siblings in the same genetic background. Values are means ±SEM for 13 hairy and 16 bald plants, means significantly different at *α*=0.01 or 0.05 are shown by * or **, respectively. b) Mean outlines of the leaves for each genotype are shown in green, with +1 SEM in grey.

## DISCUSSION

*H* encodes a member of the land plant-specific CC clade of GRXs. Other members of this clade act as adaptors between DNA-binding TGA transcription factors and transcriptional co-repressors of the TOPLESS (TPL) family (Uhrig *et al.*, 2017). As potential oxioreductases they may also catalyse reversible redox modification of target proteins (Xing *et al.*, 2005; Xing & Zachgo, 2008). Within the CC clade, the parologous ROXY1 (ROX1) and ROX2 proteins of Arabidopsis provide a precedent for developmental regulation. ROX1 binds to the TGA PERIANTHIA, reducing its ability to promote petal primordia (Li *et al.*, 2009; Murmu *et al.*, 2010; Xing *et al.*, 2005) and acts with ROX2 to repress somatic cell fate during microspore development (Xing & Zachgo, 2008), a function that predates the monocot-dicot divergence (Wang *et al.* 2009; Yang *et al.*, 2015). H shares the motifs required in other GRXs for oxioreductase activity and in ROX1 for interaction with TGAs and TPL co-repressors (Supplemental Fig. S2) and therefore might inhibit TGAs or other factors that promote trichome fate. Although no TGA is known to be involved in trichome development, identification of H-interacting proteins will allow this to be tested.

H function appears confined to part of the Lamiales, while multicellular trichomes similar in morphology to those of *Antirrhinum* are more widespread, consistent with *H* having been recruited to control an existing mechanism of trichome development. This idea is also consistent with the evidence that *H* is not necessary for the normal spacing or morphology of trichomes, which occur in the absence of H activity. In terms of its recruitment, *H* has diverged in function from its closest expressed paralogue in *Antirrhinum, GRX6c*, which is phase-regulated but not expressed in the epidermis. Several other CC-clade GRX proteins appear functionally interchangeable within species so their distinct roles likely reflect their different domains of expression (Li *et al.*, 2011; Uhrig *et al.*, 2017; Wang *et al.*, 2009). A shift in *GRX* expression to the epidermis, following gene duplication in the Lamiales, may therefore have been sufficient to bring trichome specification under control of *H*.

Evolutionary changes in trichome distribution within the genus *Antirrhinum* appear relatively unconstrained by either mutation or gene-flow. Yet, with the exception of a few polymorphic populations, distribution has remained correlated with the other morphological characters that define subsections (Wilson & Hudson, 2011) and reflect differences at multiple, unlinked loci (Costa *et al.*, 2012; Feng *et al.*, 2009; Langlade *et al.*, 2005). This is true even of *A. grossi* and *A. hispanicum*, which have evolved *Kickxiella-like* morphologies independently and carry recent *h* mutations, and *A. siculum* in subsection *Antirrhinum*, which carries a second-site suppressor of its mutant *h* allele. Such persistence and convergent evolution of character combinations suggests that the characters, which include hair distribution, are kept together by divergent selection. Though *H* may also suppress leaf growth, selection on leaf size is unlikely to have driven variation in trichomes because *Kicxkiella* species have the smallest leaves in the genus (Langlade *et al.*, 2005; Wilson & Hudson, 2011). A more plausible possibility is that a fitness cost of producing or bearing trichomes favours the bald character in the habitats to which subsections *Antirrhinum* and *Streptosepalum* are adapted or in the large, upright morphologies of these subsections. Like bald *Antirrhinum* species, many angiosperms have a restricted distribution of trichomes, often related to developmental phase (reviewed by Johnson, 1975; Poethig, 1990), consistent with a benefit from limiting them to the most vulnerable or costly parts of the plant. NILs differing only for H function could be used to examine these possibilities for *Antirrhinum.*

## MATERIALS & METHODS

### Plants and genetics

The origins of *Antirrhinum* species and the inbred *A. majus* line JI.7 are shown in Supplemental Table S3. Their taxonomy has been considered previously (Wilson & Hudson, 2011). Creation of near-isogenic lines (NILs) is described elsewhere (Costa *et al.*, 2012). A NIL carrying the *A. charidemi* allele of the H-linked *CYCLOIDEA* (*CYC*) locus were identified by a *Kpn* I CAPS after amplification with primers 5’-TCCTCCCTTCACTCTCGCGC-3’ and 5’-TGGCGCATAGCTGGTTCGAC-3’. Self-pollination of one of these NILs produced a low proportion (6%) of progeny with the hairy phenotype, suggesting that it was an *H^m^*/*h^c^* heterozygote and also heterozygous *s^m^*/*S^c^* at the self-incompatibility (*S*) locus. To obtain an *H^m^*/*h^c^* NIL lacking the active *S^c^* allele, a hairy offspring, (likely genotype *h^c^s^m^*/*h^c^S*^c^), was back-crossed to JI.7 and an individual inheriting the recombinant *h^c^s^m^* haplotype identified by a *Dpn* II CAPS in an F-box component of *S* (Lai *et al.* 2002), amplified with primers 5’-GTGCTTTCCTTCCACGATGT-3’ and 5’-CCTGGTTCAAACTGAT CAAG C-3’. A second recombination, on the opposite side of *h^c^* from *S*, occurred in this process to uncouple *h^c^* from *CYC^c^*.

### Whole-mount preparation of leaves

To examine trichome morphology, leaves were soaked in 100% ethanol for several days to remove chlorophyll, rehydrated in phosphate buffered saline (PBS) and softened and cleared in 0.5 M NaOH for 1 hour at 60°C. After washing in PBS, samples were stained in 1% safranin for 2 minutes before mounting in water.

### Leaf allometry

To test for effects of the *Hairy* introgression on leaf shape and size, bald (*n*=16) and hairy (*n*=13) progeny of the heterozygous NIL were grown together in a glasshouse and leaves from m2-13 harvest when the first flower opened. Leaves were flattened and scanned and leaf areas calculated with Fiji. Areas of leaves from each metamer were compared between genotypes with 2-tailed *t*-tests, after Shapiro-Wilk tests for normality. To compare the shapes and sizes of leaves at all nodes, each leaf outline was converted to a series of points using AAMToolbox, so that each leaf was described by 53 2D co-ordinates (Supplemental Fig. S6). For each node, the sets of points for all plants were rotated and translated to minimise variance within the dataset. Point sets for all leaves of a plant were then combined and the mean position of each point in the hairy or bald plants used as the mean shape for that genotype, and plotted with its standard error (Schindelin *et al.*, 2012).

### Quantitative RT-PCR

Three developmental stages were sampled: aerial parts of seedlings ~16 days after germination (with m2-m3 leaf primordia), shoot apices with 5 mm-long leaves at m5, and inflorescence apices. At least three tissue samples, each from multiple individuals, were processed separately. RNA was extracted with Trizol reagent, purified on Invitrogen Purelink columns and incubated with DNase to remove genomic DNA contamination, as confirmed by PCR. cDNA synthesis was primed from oligo-dT and real-time amplification from cDNA templates carried out in a Roche LightCycler 96, with at least two technical replicates of each primer-template combination. Expression was quantified relative to *A. majus Ubiquitin5* (Preston & Hileman, 2010), with amplification efficiencies for each gene estimated from a serial dilution of pooled experimental templates. The same procedure was used to test the effects of ViGS on gene expression, using RNA samples from three different plants from each treatment. Primer sequences are listed in Supplemental Information Table S4. The ratio of RNA abundance for the target gene to *Ubiquitin5* was calculated for each biological replicate and differences in expression between treatments or genotypes were detected with ANOVA and Tukey’s post-hoc test.

### Pool-seq

DNA was extracted from individual progeny of a heterozygous NIL (*H^m^/h^c^*). For each phenotype--hairy (*h^c^/h^c^* genotype) or bald (*H^c^/h^c^* or *H^c^/H^c^*)–an equal quantity of DNA from each of 86 plants was pooled. Illumina Hi-seq libraries were prepared at Earlham Institute, Norwich and sequenced to generate 100 bp paired-end reads. Reads were cleaned with Trimmomatic (Bolger *et al.*, 2014), aligned to a draft *A. majus* genome with Stampy (Lunter & Goodson, 2011) and PCR duplicates removed. A pileup file for the two pools was created from their BAM files and converted to sync format in Popoolation2 (Kofler *et al.*, 2011). Popoolation2 was used to calculate mean *F_ST_* values between the two phenotype pools for SNPs and short indels in 10 kb bins and to test significance of association between of SNPs and trichome phenotype. Because the hairy pool carries only the *h^c^* allele, while an average of one third of the alleles in bald plants will be *h^c^*, a SNP at *H* has an expected *F_ST_* of ~0.5. The order of genome sequence scaffolds along chromosomes was inferred from recombination mapping data.

The region containing *h* was delimited further by flanking recombination events, detected as *A. majus* polymorphisms in the pool of hairy phenotypes (genotype *h^c^/h^c^*). To minimise the possibility of falsely calling polymorphisms that were sequencing errors, the criterion set for detection of an *A. majus* SNP was either a single sequence feature covered by at least two independent reads or the outermost of two features within 250 bp covered by different reads. Mapping of reads to the reference was also checked manually in IGV (Robinson *et al.*, 2011) to confirm that the polymorphisms flanking *h* were not the result of inconsistent read alignment around indels.

### RNA-seq

RNA was extracted from vegetative shoot apices containing m5 primordia onwards, from multiple individuals of either the *hairy* NIL or its *A. majus* progenitor. The vegetative state of the apices was confirmed by reverse-transcription PCR of the *FLORICAULA* gene, which is expressed only after the transition to reproductive development (Bradley *et al.*, 1996), using primers 5’-GGAAGTGAGGCGGAGGCA-3’ and 5’-ACCCGCCCCCATCATTC-3’. cDNA libraries for RNA-seq were made from the RNA pools and sequenced to generate 100 bp single-end reads at Glasgow Polyomics. Reads were cleaned and trimmed as for genomic sequences, and mapped onto the draft *A. majus* genome and quantified with the Tuxedo suite (Trapnell *et al.*, 2012).

### Virus-induced gene silencing (ViGS)

Plasmids pTRV1 and pTRV2Δ2b, carrying sequences that express modified versions of the bi-partite genome of Tobacco Rattle Virus (TRV) (Liu *et al.*, 2002), were used for ViGS. To develop a reporter for silencing, 344 bp of the single-copy *Phytoene Desaturase (PDS)* gene of *A. majus* was identified by homology, amplified from cDNA and cloned into the pTRV2 vector. *PDS* and test gene sequences were fused by overlap PCR before cloning. Either the whole ORF or 237 bp of the 3’-UTR from 5 nt after the stop codon were used for *H* and the whole ORF for *GRX6c* (Supplemental Fig. S8).

Agrobacterium cells (GV3101) carrying either pTRV1 or pTRV2 were mixed and infiltrated into leaves of *Nicotiana benthamiana.* After 5-7 days leaves were ground in 1 mM phosphate buffer to obtain an infectious virus extract. Cotyledons and m2 leaves of *Antirrhinum* or *Misopates orontium* seedlings were inoculated by rubbing with the virus extract and Al_2_O_3_ abrasive. The effect of silencing *PDS* (bleaching) was visible within ~7 days of infection. The *A. majus PDS* sequence caused ViGS in *M. orontium* so was also used to monitor silencing in this species.

### *In situ* hybridization to RNA

RNA was detected in sections of fixed tissue (Rebocho *et al.*, 2017). To produce digoxigenin-labelled RNA probes, the 3’-UTR of *A. majus H* or *GRX6c* were cloned in pJET1.2, amplified as a fusion with the vector’s T7 promoter sequence and the PCR products transcribed with T7 RNA polymerase and digoxigenin-labelled dUTP. Cell walls were counterstained with calcofluor white and imaged in combined white and UV light. Because apices were too large to be included in a single image, multiple images were stitched with the photomerge function of Adobe Photoshop.

### Phylogenetic analyses

*H* sequences of *Antirrhinum* species were amplified from genomic DNA with primer pairs H-L/H-R, H-L/H-R2, or H-L3/H-R3 (Supplemental Table S4), using Q5 polymerase (NEB). For the outgroup species *Chaenorrhinum origanifolium* and *M. orontium*, they were amplified from cDNA of vegetative shoot apices by 5’ and 3’ RACE (Scotto-Lavino *et al.*, 2006a; b) using primers GRX-CR-R, or GRX-RACE-R3 and RACE adaptor primers. Direct sequencing of PCR products indicated heterozygosity in most *Antirrhinum* accessions. Therefore products of an independent PCR were cloned in pJET1.2 and clones sequenced until both haplotypes had been recovered from a heterozygous individual. PCR-generated mutations were eliminated by comparing sequence traces from clones and initial PCR products.

The *H* coding sequences were aligned with MUSCLE in MEGA (Edgar, 2004). MEGA was also used for maximum likelihood (ML) estimates of phylogeny, with *M. orontium* as outgroup, use of sites present in ≥75% of sequences and a model with HKY substitution and gamma distribution of rates, as chosen by jModeltest2 (Darriba *et al.*, 2012). Bootstraps percentages are from 500 replicates.

To examine the likely origin of H, similar proteins were identified by blast searches of the inferred proteomes of *Arabidopsis thaliana*, Mimulus *(Erythranthe guttata)* and tomato (Supplemental Table S5). Other *A. majus* proteins were inferred from a combination of genome and cDNA sequences and *Misopates* and *Chaenorrhinum* proteins from amplified cDNAs.

Partial amino acid sequences (residues 11-107 of *A. majus* H) were aligned with MUSCLE. The most likely ML tree was identified in MEGA using a JTT model with gamma distributed rates, as suggested by Prottest2.4 (Abascal *et al*., 2005), and tested with 500 bootstrap replicates. The tree was rooted with reference to a larger tree (Supplemental Fig. S7), produced using RaxML (Stamatakis, 2014) under a similar model.

## AKNOWLEDGEMENTS

YT and MB were supported by studentships from Darwin Trust of Edinburgh and the Biotechnology and Biological Sciences Research Council (BBSRC; grant BB/J01446X/1), respectively. We thank Yongbiao Xue for the *A. majus* reference sequence, Annabel Whibbley and Enrico Coen for *A. sempervirens* and *A. pseudomajus* resequencing data, Yana Aleksandrova for initial mapping studies, Fei Yue and Douglas Pyott for advice on ViGS and Jill Harrison and Justin Goodrich for comments on the manuscript.

