## Supplemental Information for "Loss of a glutaredoxin gene underlies parallel evolution of trichome pattern in *Antirrhinum*"

**SUPPLEMENTAL FILES, Tan *et al.***

***Allelism tests with A. siculum***

Crosses of the $H^m/h^c$ NIL to bald *A. siculum*, in either direction, produced very few viable seeds–the only seedling obtained proved to be hairy and to have had inherited the $h^c$ allele. A single F1 progeny of the cross to *A. majus* ($H^m/H^m$) was bald. *A. siculum* showed higher fertility when crossed with *A. molle* (hairy phenotype, *h/h* genotype) and all progeny in this case were hairy, suggesting that *A. siculum* lacked H activity. An F2 population produced by selfing the $h^{siculum}/h^c$ F1 from the NIL segregated hairy and bald plants in a ratio that was not significantly different from 3:1 (476 hairy, 132 bald; $\chi^2$ $p$=0.06). Baldness in *A. siculum* can therefore be explained by a lack activity of a second locus, *Bald* (*B*), required for formation of the trichomes that are suppressed by *H*. In this case *A. siculum* would have genotype *h/h b/b, A. molle h/h B/B*, the NIL *H/h B/B*, and *A. majus H/H B/B*. The F1 offspring of the cross to *A. molle* would all inherit the dominant *B* allele (permissive for hair formation) and, being homozygous *h/h*, produce hairs. The *h/h B/b* F1 plant produced by crossing to the NIL carries a functional *B* allele and is hairy, while around one quarter of its F2 progeny are *b/b* homozygotes and bald.

**Estimating map distance between *Hairy* and the *S locus***

To estimate the distance between *H* and *S*, *A. majus* was crossed with *A. charidemi* to give an F1 plant with genotype $H^ms^m/h^cS^c$, which was self-pollinated. It was assumed to have produced recombinant gametes ($H^mS^c$ and $h^c/s^m$) each with a frequency of $\frac{r}{2}$, where *r* is the recombination fraction, and non-recombinant gametes ($H^ms^m$ and $h^cS^c$) each with a frequency of $\frac{(1-r)}{2}$. However, because pollen carrying $S^c$ is rejected on self-pollination, F2 plants inherit only the parental $H^ms^m$ or recombinant $h^cs^m$ haplotype through pollen, with frequencies of 1-*r* and *r*, respectively. Of the possible F2 genotypes, two are $h^c/h^c$ homozygotes with hairy phenotypes: $h^cs^c/h^cS^m$, occurring at a frequency of $r\frac{1-r}{2}$, and $h^cs^m/h^cs^m$ with a frequency of $r\frac{r}{2}$. The predicted proportion of hairy progeny is therefore $r\frac{1-r}{2}+r\frac{r}{2}=\frac{r}{2}$, and of bald progeny $1-\frac{r}{2}$, so the expected proportion of F2 progeny that are hairy is $\frac{r}{2-r}$. The observed proportion was $\frac{18}{267}$, therefore $r$≈0.126. Applying Kosambi's mapping function this is equivalent to a map distance of 12.9 cM.

| **Trait** | **Hairy** ($H^m/H^m$) *n*=13 | **Bald** ($h^c/h^c$) *n*=16 | ***p*-value‡** |
|---|---|---|---|
| First spiral node | 11.8 (±0.4) | 12.4 (±0.6) | 0.37 |
| First flowering metamer* | 17.8 (±1.4) | 19.8 (±1.0) | 0.25 |
| Metamer of longest leaf† | 11.0 (±0.9) | 10.6 (±0.5) | 0.66 |

**Table S1 Traits related to developmental phase in hairy and bald NILs**

Values are means for hairy plants or bald plants in a near-isogenic background ± their standard errors. *: first flowering metamer is equivalent to the number of metamers that produced leaves before flowering and is therefore a proxy for flowering time; †: leaf length initially increases with increasing metamer number then declines; the position of the longest leaf is therefore a proxy for heteroblastic variation; ‡ probabilities from Students *t*-tests of means being the same.

| Male parent | Subsection* | Parent phenotype | N bald F1 | N hairy F1 | *p*(1:1)† |
|---|---|---|---|---|---|
| *A. majus* | *A* | Bald | 68 | 0 | - |
| *A. linkianum* | *A* | Bald | 30 | 0 | - |
| *A. pseudomajus* | *A* | Bald | 31 | 0 | - |
| *A. striatum* | *A* | Bald | 37 | 0 | - |
| *A. tortuosum* | *A* | Bald | 63 | 0 | - |
| *A. siculum* | *A* | Bald | 0 | 1‡ | - |
| *A. hispanicum* | *K* | Hairy | 17 | 14 | 0.59 |
| *A. boissieri* | *K* | Hairy | 16 | 21 | 0.41 |
| *A. graniticum* | *A* | Hairy | 20 | 22 | 0.76 |
| *A. lopesianum* | *K* | Hairy | 6 | 6 | 1.00 |
| *A. molle* | *K* | Hairy | 19 | 15 | 0.49 |
| *A. rupestre* | *K* | Hairy | 9 | 11 | 0.65 |
| *A. pulverulentum* | *K* | Hairy | 27 | 22 | 0.48 |
| *A. subbaeticum* | *K* | Hairy | 7 | 6 | 0.78 |
| *A. valentinum* | *K* | Hairy | 19 | 17 | 0.74 |

**Table S2 Allelism tests**

Each species was crossed as pollen parent to the NIL that was heterozygous $H^m/h^c$ and the numbers of hairy and bald phenotypes in the progeny recorded. *: *A*, subsection *Antirrhinum*, *K*, subsection *Kickxiella*; † *p*-values from $\chi^2$-tests fitting the observed numbers of F1 phenotypes to the 1:1 ratio expected if the hairy parent was homozygous *h*/*h*; ‡ few viable seeds were produced by crossing *A. siculum* to the NIL.

| | | | | |
|---|---|---|---|---|
| *Antirrhinum* | *A. australe* Rothm. | L084* | El Boyar, Cádiz, Spain | *xxvi* |
| | | L091* | Gaucín, Málaga, Spain | *xiv, xxvi* |
| | | L095* | El Burgo, Málaga, Spain | *x, xii, xiii, xiv, xxiii* |
| | *A. barrelieri* Boreau | L150 | Cádiar, Granada, Spain | *vii* |
| | *"A. barrelieri"* | L167 | Taineste, Morocco | *xiv* |
| | *A. graniticum* Rothm. | L040* | Bragança, Bragança, Portugal | *iv, v, vii, xiv, xvii* |
| | | L069 | Ledanca, Guadalajara, Spain | *vii* |
| | | L116* | Celorico da Beira, Guarda, Portugal | *xviii* |
| | *A. latifolium* Miller | AC1066 | St Martin d'Entraunes, Alpes-Maritimes, Fra. | *xvi* |
| | *A. cirrhigerum* Filcaho | L114 | Praia de Mira, Coimbra, Portugal | *viii* |
| | *A. linkianum* Boiss & Reuter | L108 | Almada, Setúbal. Portugal | *xiv* |
| | *A. litigiosum* Pau | L003 | Cheste, Valencia, Spain | *iii* |
| | *A. majus* L. | | JI.7 from John Innes Centre | *vii* |
| | *A. pseudomajus* Rouy | L053 | Minerve, Hérault, France | *ix* |
| | *A. striatum* Rothm. | AC1125 | Alet-les-Bains, Aude, France | *i* |
| | *A .tortuosum* Rouy | L092 | Casares, Málaga, Spain | *xiv* |
| | *A. siculum* Millar | AC1177 | Taormina, Sicily, Italy | *xxviii* |
| *Kickxiella* | *A. boissieri* Rothm. | L018 | Guadahortuna, Granada, Spain | *xv, xxii* |
| | | L104 | Ermita Virgen de la Sierra, Cordoba, Spain | *xix* |
| | *A. charidemi* Lange | E023 | Cabo de Gata, Almería, Spain | *xx* |
| | *A. grosii* Font Quer | L175 | Sierra de Gredos, Toledo, Spain | *xi* |
| | *A. hispanicum* Chav. | L099 | Almargen, Málaga, Spain | *ii* |
| | *A. lopesianum* Rothm. | L038 | Vimioso, Bragança, Portugal | *xxiv* |
| | *A. microphyllum* Rothm. | L072 | Sacedón, Guadalajara, Spain | *xxx* |
| | | L073 | Pantano de Buendía, Guadalajara, Spain | Δ† |
| | | L074 | Buendía, Guadalajara, Spain | *xxix* |
| | *A. molle* Lange | E051 | Gerri de la Sal, Lleida, Spain | *xxv* |
| | *A. mollisimum* Rothm. | L021 | Enix, Almería, Spain | *xvii* |
| | *A. pertegasii* Rothm. | E065 | Castellón de la Plana, Castellón, Spain | Δ |
| | *A. pulverulentum* Lazaro | L068 | Pelegrina, Guadalajara, Spain | Δ |
| | | L070 | Alcorlo, Guadalajara, Spain | *xxvii* |
| | | L077 | Poveda de la Sierra, Guadalajara, Spain | Δ |
| | *A. rupestre* Rothm. | L139 | Capileira, Granada, Spain | *xxi* |
| | *A. sempervirens* Lapeyre | L050 | Col d'Aubisque, Hautes-Pyrenees, France | Δ |
| | | L052 | Luz-Saint-Sauveur, Hautes-Pyrenees, France | Δ |
| | *A. subbaeticum* Guemes | E72 | Albacete, Albacete, Spain | Δ |
| | *A. valentinum* Font Quer | AC1173 | La Drova, Valencia, Spain | Δ |
| *Streptosepalum* | *A. meonanthum* Hoffmans & Link | L118 | Manteigas, Guarda, Portugal | *xi, xiv* |
| | *A. braun-blanquetii* Rothm. | E20 | St Pietro de Villanueva, Asturias, Spain | *xiv* |
| Other species | *Chaenorhinum origanifolium* Fourr. | | Gijón, Asturias, Spain | *CoGRX* |
| | *Misopates orontium* Raf. | | Unknown | *MoHairy* |

**Table S3 Accessions of *Antirrhinum* and related species** *: polymorphic population with both bald and hairy individuals, †: Representatives of endemic *Kickiella* species in which the *H* locus could not be detected and is presumed to have been deleted are shown with Δ. For relationships of haplotypes (Roman numerals) please see Supplemental Fig. S9.

| Use | Gene | Primer Name | Sequence (5'-3') |
|---|---|---|---|
| qPCR | *A. majus H* | AmHQ-F | TCTTGTCTTTTCCACCTGTCA |
| | | AmHQ-R | TGAATATCACCACCGGATTCTC |
| | *A. majus GRX6c* | AmGRX6cQ-F | TGGCTCTAGTTCCTAAGGAGAA |
| | | AmGRX6cQ-R | CACAAGCCTACAGAGCTACTAATC |
| | *A. majus PDS* | AmPDSQ-F | TCTTTGTAATGGACGGCAAG |
| | | AmPDSQ-R | ACTTGCCAAACTCTTCCCTG |
| | *A. majus Ubi5* | AmUbiQ-F | CCGAACCATCAGACAAACAAAC |
| | | AmUbiQ-R | TACCCTGGCCGACTACAATA |
| Phylogenetic analysis | *Antirrhinum H* | AmH-R | GTAGTCCTATACAAATTAATACGTA |
| | | AmH-F | ACAGAGTATACGCCTCGAT |
| | | AmH-F3 | GTTTCCCTGGAATCAACCAC |
| | | AmH-R3 | AACACCGTCGCTGTTGCTC |
| | | AmH-R2 | ACAGTCCTATACAAATTAATATG |
| | *Misopates Hairy* & *Chaenorrhinum GRX* | GRX-RACE-R3 | ACCAAGTGGCGTACGAAATTA |
| | | GRX-CR-R | GAATCCGGTGGTGATATTCA |

**Table S4 PCR primers**

| | |
|---|---|
| ***Arabidopsis thaliana* TAIR10** | |
| AtROXY1 | At3g02000 |
| AtROXY2 | At5g14070 |
| ***Mimulus guttatus* v2.0** | |
| MigF00038 | Migut.F00038 |
| MigK01018 | Migut.K01018 |
| MigK01479 | Migut.K01479 |
| MigM01237 | Migut.M01237 |
| ***Solanum esculentum* iTAG2.4** | |
| S01g009890 | Soly01g009890 |
| S01g009900 | Soly01g009900 |
| S01g009910 | Soly01g009910 |
| S05g015910 | Soly05g015910 |

**Table S5 Origins of GRX peptide sequences**

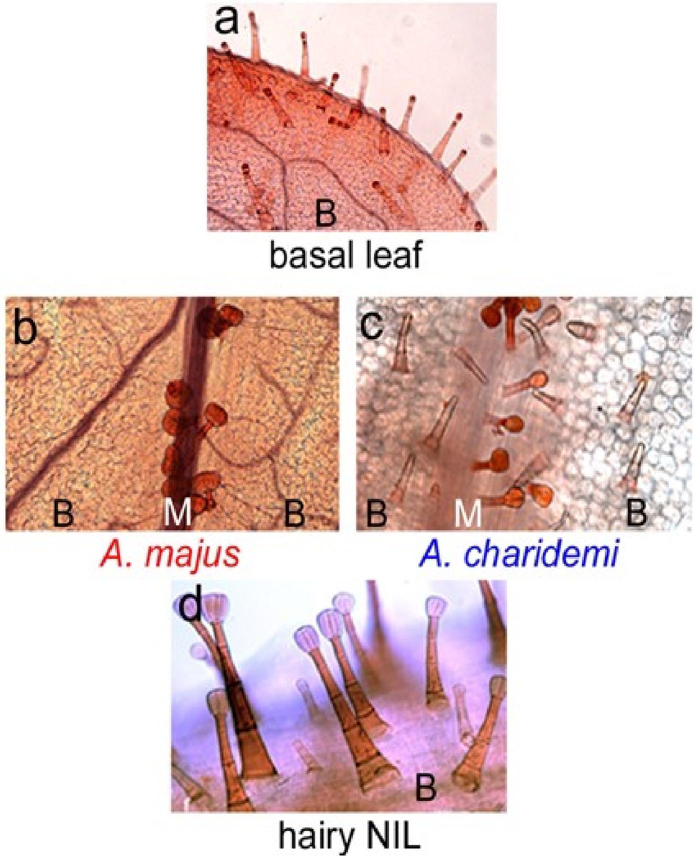


**Figure S1 *Hairy* underlies variation in trichome distribution but not trichome morphology**

a) Both *A. majus* and *A. charidemi* produce glandular secretory trichomes from the blade (B) and midrib (M) of leaves below metamer 5 (m5). From m5 onwards *A. majus* leaves (b) make trichomes only from the adaxial midrib (M), and leaf blades are bald. At the same nodes, *A. charidemi* (c) has the same type of glandular trichomes on the midrib as *A. majus*, however it produces short, non-secretory trichomes from the leaf blade. The hairy NIL, homozygous for the *A. charidemi* $h^c$ allele, produces trichomes on the blades of leaves from m5 onwards (d), as does its *A. charidemi* parent. However, these trichomes are glandular, unlike those of *A. charidemi*, indicating that the *Hairy* gene underlies variation in trichome distribution, but not trichome morphology.

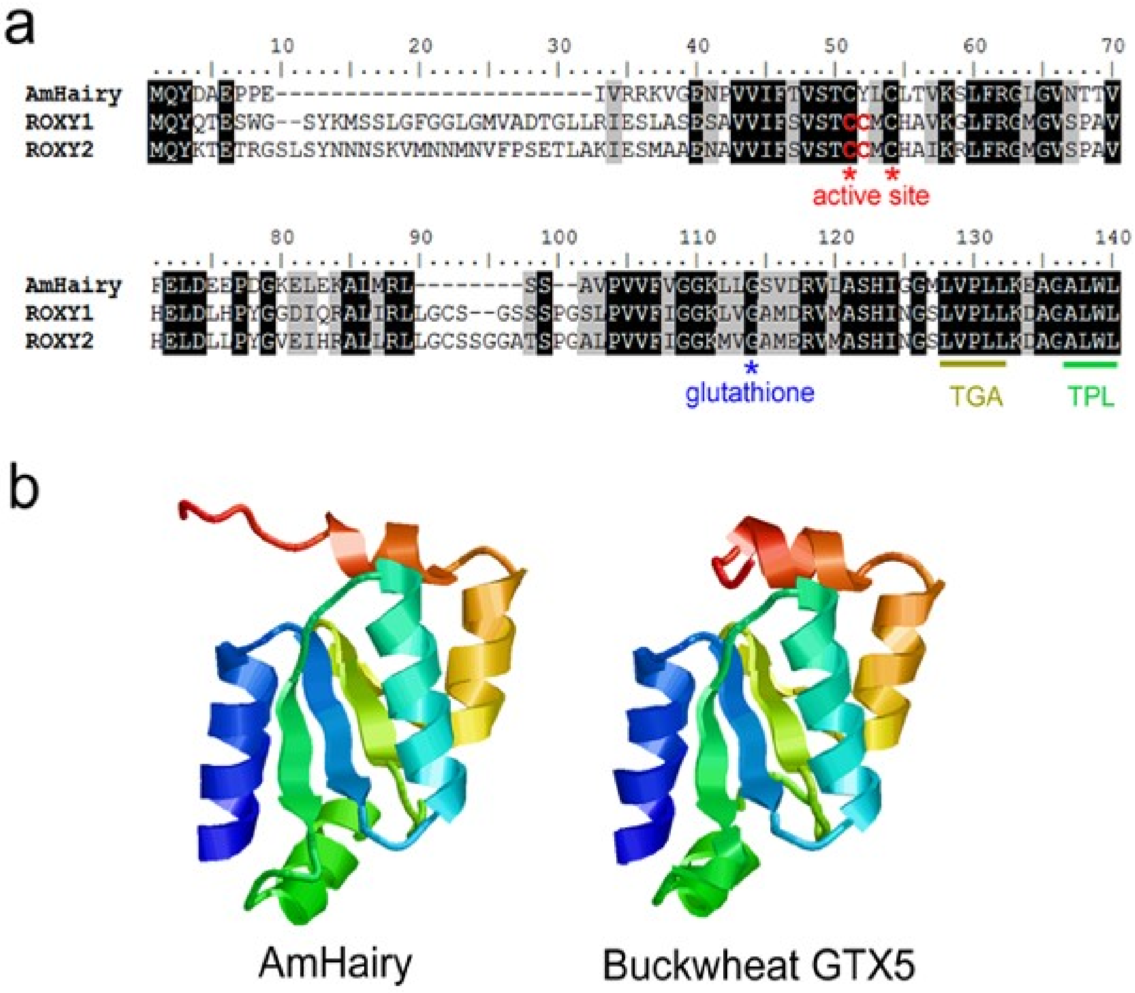


**Figure S2 *Hairy* encodes a CC-family glutaredoxin**

a) Clustal alignment of the predicted full-length Hairy protein from *A. majus* with the most similar proteins from *Arabidopsis thaliana*–ROXY1 (ROX1, the product of At3g02000) and ROX2 (At5g14070). Identical residues are boxed in black, conservative substitutions in grey. The two catalytically active cysteines, conserved among almost all other CC-family glutaredoxins, and the glycine residue required for glutathione cofactor binding, are highlighted. Although Hairy is nested within the CC-family (Supplemental Fig. S7), it lacks the second of the two adjacent active-centre cysteins (CC in red type), which are ancestral in the family, retained by most other members (1), and give the family its name. The hydrophobic L**LL motif, required by ROX1 and ROX2 for interaction with TGA transcription factors (2), and the ALWL motif required for interaction with transcriptional co-repressors of the TPL family (3) are also shown. All the motifs are also present in the *H* paralogue *GRX6c* (not shown). b) Homology-informed secondary structure prediction for *A. majus* Hairy, made with Phyre2 (4), compared to the experimentally-determined structure of a GRX from buckwheat, expressed in *E. coli* (5).

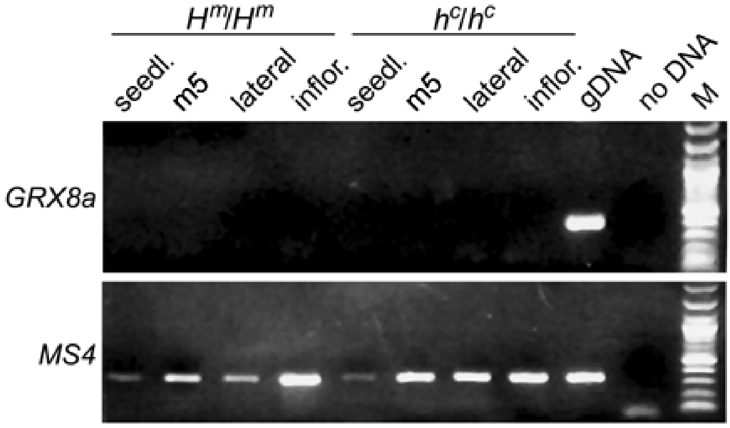


**Figure S3 Testing expression of the *H* paralogue *GRX8a***

cDNA was made from different aerial tissues of near-isogenic bald plants ($H^m/H^m$), homozygous for *A. majus* alleles of both *H* and its closely-linked paralogue *GRX8a*, or hairy plants ($h^c/h^c$), homozygous for *A. charidemi* alleles at both loci. Expression of *GRX8a* could not be detected by RT-PCR using primers within its single predicted exon, in either the presence or absence of H activity, though a genomic DNA (gDNA) template could be amplified under the same conditions. cDNA from the housekeeping gene *Methionine synthase 4* (*MS4*) was detected in all samples. Tissues were aerial parts of whole seedlings (seedl.), vegetative shoot tips with 5 mm leaves at metamer 5 (m5), tips of vegetative axillary shoots from above m5 (lateral) and inflorescence apices (infl.). M is a size marker.

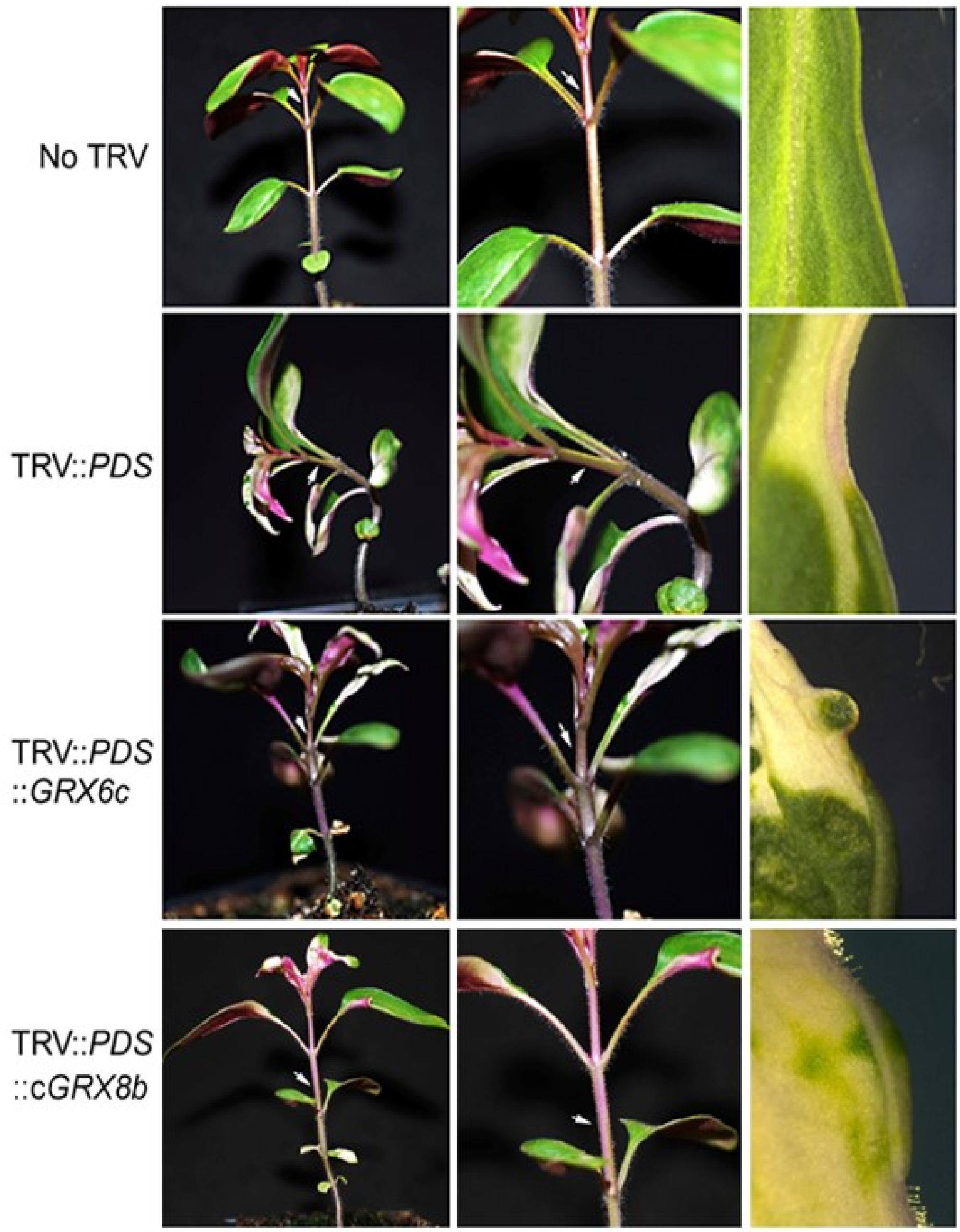


**Figure S4 ViGS of the *H* paralogue *GRX6c***

ViGS using a parts of the *GRX6c* and *PDS* sequences reduced *GRX6c* RNA abundance to below 20% of normal levels (Fig. 3f), but did not affect trichome development in bleached tissues showing ViGS of *PDS* (third row). In contrast, ViGS with part of the coding sequence of *GRX8b* (bottom row) allowed formation of ectopic trichomes. The right-hand panels show parts of representative metamer 5 leaves. Arrows show the position of the last stem trichome, which is unaffected by reduced *GRX6c* expression.

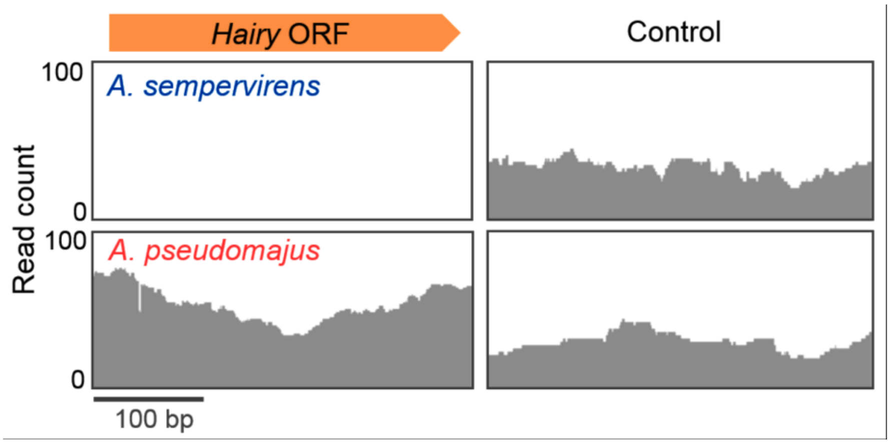


**Figure S5 Deletion of *H* in a group of endemic *Kickxiella* species**

a) Coverage of genome resequencing reads at *A. majus Hairy* (left) or an unlinked single-copy region (right). The reads are from *A. sempervirens*, one of the endemic subsection *Kickxiella* species from which *H* could not be amplified, or *A. pseudomajus* (genotype *H*/*H* from subsection *Antirrhinum*). Reads were mapped with the same parameters.

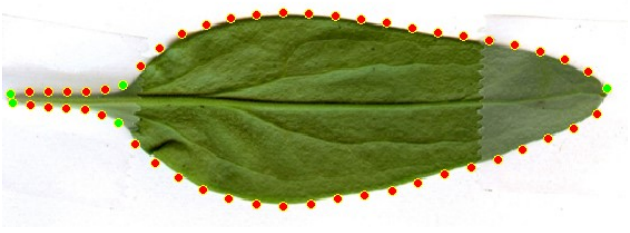

**Figure S6 Describing leaf outlines numerically**

Fifty-three points were positioned around the outline of each leaf. The green points were placed manually, the red points spaced automatically at equal distances between them.

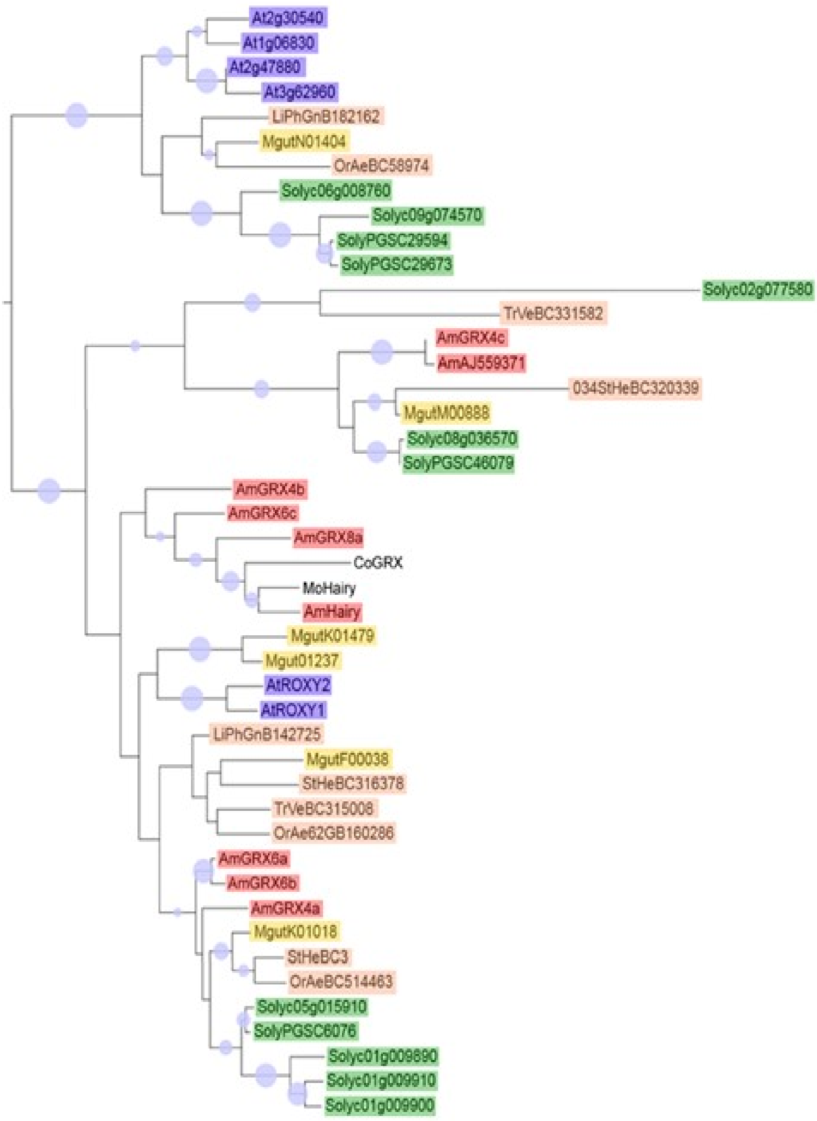


**Figure S7 Inferred relationships of Hairy-like GRX proteins**

The most probably ML tree was obtained using RaxML 8.0 with parameters similar to those used for the subclade containing Hairy (Fig. 5a). Sequences from the Orobanchaceae species *Orobanche aegyptiaca* (OrAe), *Striga hermontheca* (StHe), *Triphysaria versicolor* (TrVe) and *Lindenbergia philippensis* (LiPh), which are coloured carmine, were translated from EST databases of the Parasitic Plant Genome Project (http://ppgp.huck.psu.edu/). They were not used for the Hairy subtree in Figure 5a because of the possibility of other paralogues remaining unsampled as ESTs. The remaining sequences from *A. majus* (red), Mimulus (yellow), tomato (green) and Arabidopsis (blue) are from annotated reference genomes (Table S5). Nodes recovered in ≥50% of 520 bootstrap replicates have circles of a size proportional to the level of support.

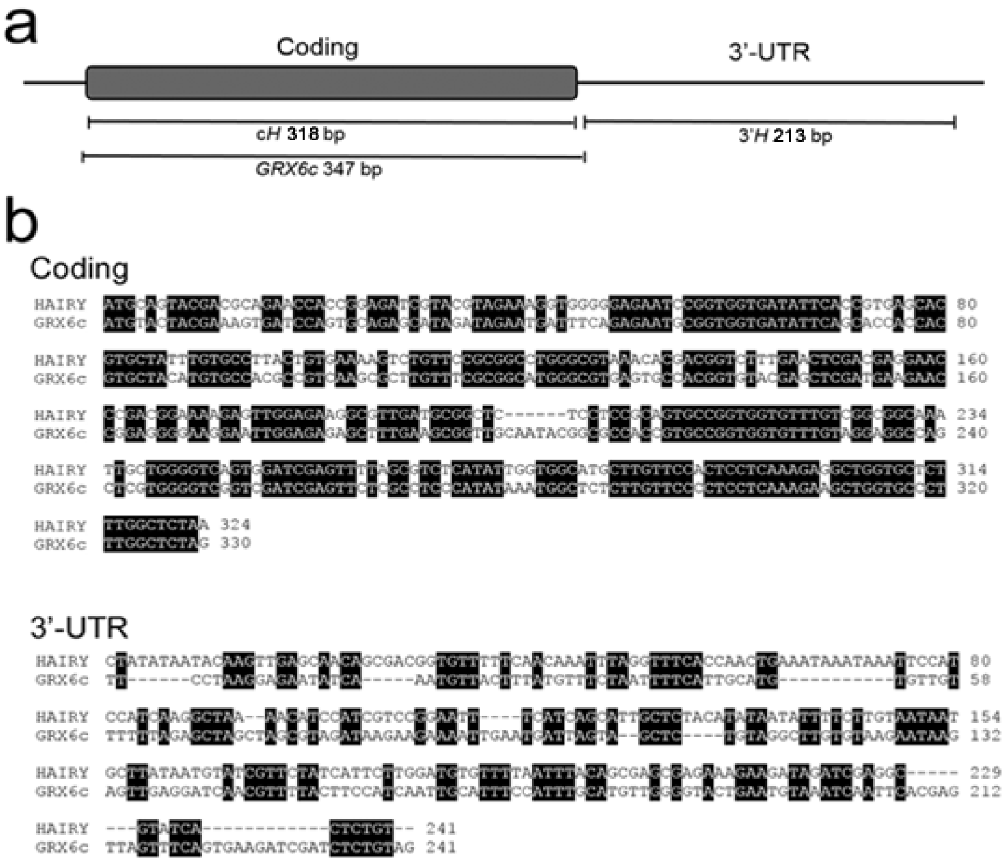


**Figure S8 *GRX* gene sequences used for ViGS**

a) ViGS of *H* was achieved either with the whole coding region (c*H*) or most of the 3'-UTR (3'*H*), attempted ViGS of the *H* paralogue *GRX6c* used a sequence extending from 2 bp upstream of the initiation codon to 38 bp downstream of the Stop codon. Both genes lack introns. b) Sequence alignments of the coding or 3'-UTRs of *H* and *GRX6c*. The longest block of identical sequence is 17 bp in length.

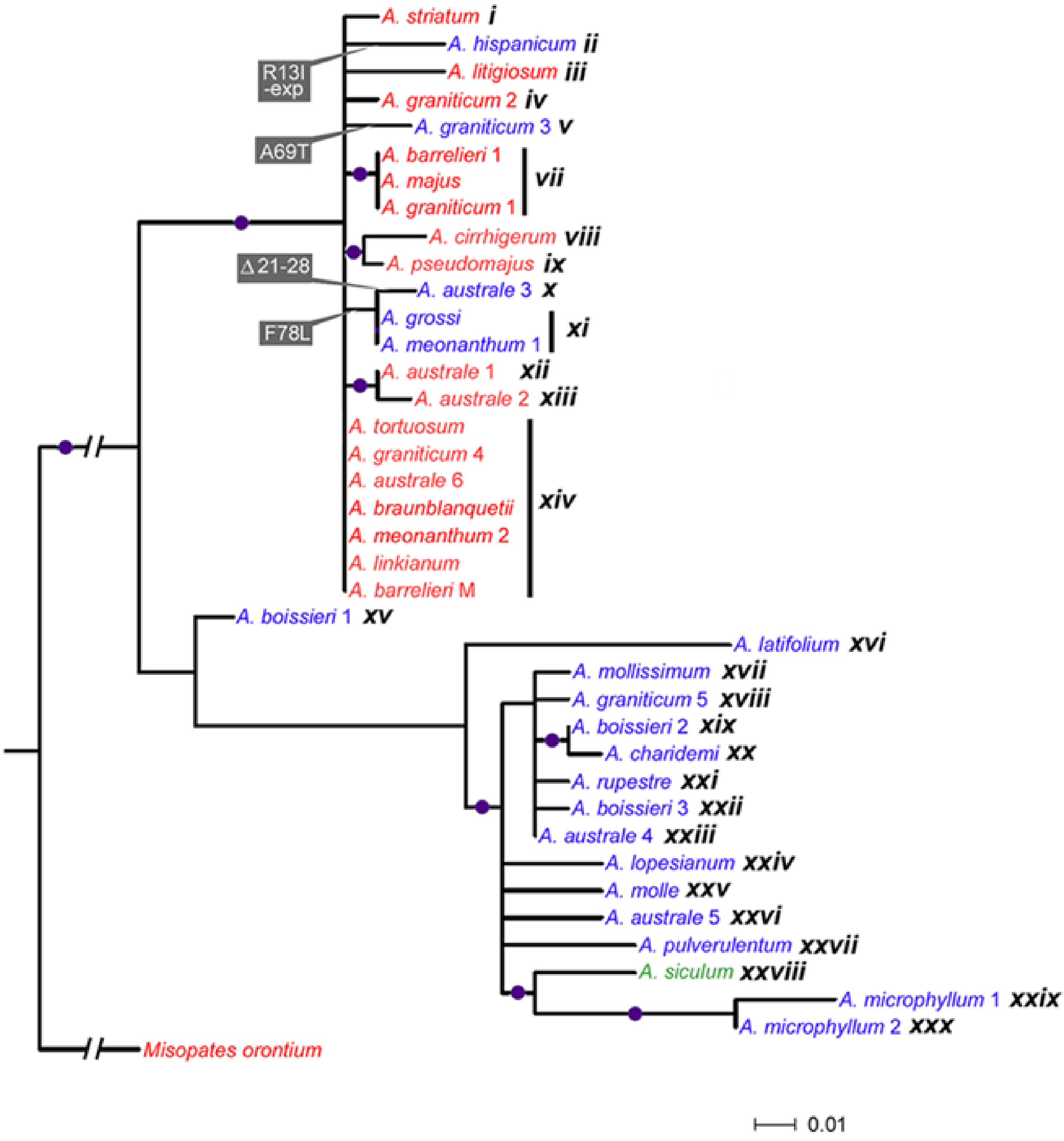


**Figure S9 Distribution of *Hairy* haplotypes among species**

*Hairy* coding sequences are named according to the species from which they originated, with Arabic numerals used to distinguish different sequences from the same species (e.g., *A. microphyllum* 1 and *A. microphyllum* 2). Because more than one species could share the same sequence, each unique sequence (haplotype) was given a number in Roman numerals (*e.g.*, haplotype *xi* was found in both *A. grossi* and *A. meonanthum*). The distribution of sampled haplotypes among species and populations is summarised in Supplemental Table S3.
